# Proteomic characterization of *Mycobacterium tuberculosis* subjected to carbon starvation

**DOI:** 10.1101/2024.11.12.623260

**Authors:** Kaylyn L. Devlin, Damon T. Leach, Kelly G. Stratton, Gyanu Lamichhane, Vivian S. Lin, Kimberly E. Beatty

**Author notes:** Corresponding author: Kimberly E. Beatty.

## Abstract

*Mycobacterium tuberculosis* (*Mtb*) is the causative agent of tuberculosis (TB), the leading cause of infectious-disease related deaths worldwide. TB infections present as a spectrum from active to latent disease. In the human host, *Mtb* faces hostile environments, such as nutrient deprivation, hypoxia, and low pH. Under these conditions, *Mtb* can enter a dormant, but viable, state characterized by a lack of cell replication and increased resistance to antibiotics. These dormant *Mtb* pose a major challenge to curing infections and eradicating TB globally. In the current study, we subjected *Mtb* to carbon starvation (CS), a culture condition that induces growth stasis and mimics nutrient-starved conditions associated with dormancy *in vivo*. We provide a detailed analysis of the proteome in CS compared to replicating samples. We observed extensive proteomic reprogramming, with 36% of identified proteins significantly altered in CS. Many enzymes involved in oxidative phosphorylation and lipid metabolism were retained or upregulated in CS. The cell wall biosynthetic machinery was present in CS, although numerous changes in the abundance of peptidoglycan, arabinogalactan, and mycolic acid biosynthetic enzymes likely result in pronounced remodeling of the cell wall. Many clinically approved anti-TB drugs target cell wall biosynthesis, and we found that these enzymes were largely retained in CS. Lastly, we compared our results to those of other dormancy models and propose that CS produces a physiologically-distinct state of stasis compared to hypoxia in *Mtb*.

## Introduction

Tuberculosis (TB) is the deadliest infectious disease in human history. *Mycobacterium tuberculosis* (*Mtb*), the underlying bacterial pathogen, causes over 10 million infections and over 1.2 million deaths per year. After more than a decade of slow decline in TB incidence, global case numbers started to increase in 2020 with the onset of the COVID19 pandemic[1]. Unfortunately, the international effort to end the TB pandemic by 2030 remains uncertain and challenging[2].

Eradication is hampered by the unusual ability of *Mtb* to survive within the host indefinitely as a latent TB infection (LTBI)[3]. An estimated 25% of the global population has a LTBI, with the potential to progress to an active infection at a later time[1]. There is no single biomarker that defines dormant *Mtb*, making it challenging to identify among the millions of latently-infected people who would most benefit from preventative treatments. Additionally, patients with active TB harbor physiologically-distinct subpopulations of *Mtb* across a spectrum from dormant to active [4], highlighting that infections are complex and heterogeneous. Dormant bacteria, or *Mtb* in a non-replicating persistent state, are challenging to kill in the host because they are phenotypically drug resistant[5].

There are a variety of *in vitro* models that attempt to replicate conditions in the host that induce *Mtb* dormancy[5, 6]. Bacteria are subjected to hypoxia, nutrient deprivation, low pH, reactive nitrogen species, alternative carbon sources (butyrate or host lipids), or a combination of these stressors. A feature of most models is that *Mtb* enters stasis, where growth is negligible over time. Upon entering stasis, *Mtb* undergoes widespread transcriptional and proteomic changes leading to reorganization of many cellular processes[7, 8]. *Mtb* slows metabolism and becomes phenotypically resistant to most drugs[9]. The most common dormancy model, the Wayne model, uses gradually-induced hypoxia to shift *Mtb* into two distinct stages of non-replicating persistence [10]. This model is well characterized [11–14]. The nutrient starvation phenotype was first described in 1933 by Loebel[15] and later developed and characterized by Betts et al. [16]. The Betts model cultures *Mtb* in carbon- and lipid-free buffer for six weeks. A closely related carbon starvation (CS) model was described by Grant and coworkers[17]. Multi-stress models have also been described[18, 19]. Occasionally, stationary phase is used as a surrogate for dormancy [20, 21]. Choice of model is usually based on the aspect of LTBI that the researcher wishes to study.

No single *in vitro* model is likely to recapitulate all features of dormant *Mtb* found in a host environment, especially since microenvironments vary within a single host[4]. However, these models remain informative by allowing us to study the effects of known host environments on *Mtb* biology and drug response[22]. There have been a variety of proteomic and transcriptomic studies on *Mtb* dormancy models, as recently reviewed[7]. However, nearly all have focused on dormancy induced by hypoxia. Nutrient deprivation is likely a pivotal stress for *Mtb in vivo*, particularly within the granuloma. The CS model captures many observed features of *Mtb* isolated from patients, including loss of acid-fast staining, drug tolerance, and low respiration[23]. CS is also easier to implement in the lab and more reproducible for drug screening compared to hypoxia[17].

Before now, there was limited information on proteomic changes associated with nutrient starvation[16, 24]. The most recent study was published over a decade ago and focused solely on secreted proteins[24]. Therefore, we sought to characterize proteomic changes in *Mtb* under CS conditions. We identified specific proteins and their abundance changes between replicating and CS conditions. Our results provide a comprehensive overview of protein-level adaptations used by *Mtb* to survive carbon starvation. We focus our analysis on changes in metabolism, cell wall biosynthesis, and drug targets.

## Experimental Procedures

### Experimental Design and Statistical Rationale

A total of 12 samples were analyzed in this study, comprised of six biological replicates in two distinctly cultured groups. A single bacterial stock was cultured and washed before splitting into two groups: the control group (replicating; n = 6) and the experimental group (carbon starvation; n = 6). The number of replicates per group was determined for sufficient power in quantitation and statistical analyses, and to ensure the reproducibility of our results (see [25]). Samples were analyzed by liquid chromatography tandem mass spectrometry in a randomized order using a random list generator.

### Mycobacterial Culture Conditions

*Mycobacterium tuberculosis* (*Mtb*) mc^2^6020 (Δ*lysA*, Δ*panCD*)[26], a double auxotrophic mutant derived from the laboratory strain *Mtb* H37Rv, was obtained from W. Jacobs’ laboratory (Albert Einstein College of Medicine and HHMI). It was handled as a BSL-2 pathogen under OHSU-approved biosafety protocols. All bacterial manipulation was done within a biosafety cabinet to minimize the potential for exposure to the pathogen.

Bacteria were thawed from frozen stocks stored at −80 °C in 30% glycerol. *Mtb* was cultured in 7H9/OADC-KPC medium (7H9 broth [BD Difco], 0.5% glycerol [Fisher, molecular biology grade], 0.05% Tween 80 [Sigma], 10% OADC [BD Difco], 80 μg/mL lysine [K, Sigma], 24 μg/mL pantothenate [P, Sigma], 0.2% casamino acids [Gibco]). Cultures were grown at 37 °C with 100 rpm in aerated polycarbonate shake flasks with a 0.2 μm filter cap (Weaton #WPFPC0500S).

Culture conditions to induce dormancy via carbon starvation (CS) were adapted from previously reported methods[17, 27]. *Mtb* was grown in 7H9/OADC-KPC to an OD_600_ of 0.8 - 1.2. Cells were washed twice with PBS and diluted to an OD_600_ of 0.2 in carbon starvation medium (7H9/Tx-KP; 7H9 broth, 0.05% tyloxapol [Sigma], 80 μg/mL lysine, 24 μg/mL pantothenate). Cultures (n = 6, 300 mL) were grown standing at 37 °C in 1 L plug-sealed bottles (Corning #430195) for 5 weeks (**Table S1**).

Matched replicating cultures were simultaneously prepared from the same washed cell stock. Cells were diluted to an OD_600_ of 0.2 in 7H9/OADC-KPC medium. Cultures (n = 6, 200 mL) were grown shaking (100 rpm, 37 °C) in aerated 500 mL shake flasks until an OD_600_ of ∼1.0 was reached. Cells were harvested through centrifugation (5 min, 4000 xg, 4 °C), washed twice with PBS, and stored at −30 °C in PBS until lysis.

### Lysate Preparation

Cells were lysed following a protocol adapted from previously described methods[28]. Frozen cell pellets were thawed on ice. Cells were lysed in PBS by mechanical disruption with a MiniLys Beadbeater (Bertin Technologies) at 5,000 rpm (3x 45 s, cooling between on ice for 180 s) using 0.1 mm zirconia/silica beads (BioSpec Products). Beads and cell debris were pelleted by centrifugation (18,000 xg, 5 min, 4 °C). After the first clarified supernatant was collected, 1 mL of 1% n-dodecyl-D-β-maltoside (Chem-Impex #21950, CAS 69227-93-6) in PBS (PBS-DM) was added to the remaining beads and cell debris. The slurry was resuspended and incubated for 30 min on ice. The second supernatant was collected after centrifugation and added to the clarified supernatant to give lysate in buffer with a final concentration of 0.5% n-dodecyl-D-β-maltoside. Total lysates were centrifuged a final time to remove residual beads (18,000 xg, 5 min, 4 °C). Lysates were filtered twice through 0.2 µm PVDF membrane filters (13 mm, Pall) to sterilize. The second filtration was done in a clean and sterilized biosafety cabinet. A bicinchoninic acid (BCA) assay (Pierce) was used to quantify the total protein concentration of all lysates.

### Mass Spectrometry Sample Preparation

Fresh lysate proteins (150 μg) were digested with proteomics-grade trypsin (Promega) overnight at RT with end-over-end rotation. Solid Phase Extraction (SPE) C18 columns (50 mg bed wt., Millipore-Sigma Supelco Discovery DSC-18) were conditioned with methanol (3 mL, under vacuum, Fisher) and rinsed with acidified water (2 mL, 0.1% trifluoroacetic acid [TFA], Fisher). The peptides from the digestion were applied to the SPE columns and slowly allowed to pass through the column (≤ 1 mL/min). Samples were washed with 4 mL of 95:5 H_2_O:acetonitrile (ACN, Fisher) with 0.1% TFA. Columns were allowed to go to dryness, and column tips were wiped to remove any residue. Peptides were slowly eluted from columns with 20:80 H_2_O:ACN with 0.1% TFA (1 mL), under vacuum. The samples were dried in a SpeedVac concentrator and resuspended in HPLC-grade water (50 µL). Peptide concentrations were quantified via BCA assay. Samples were normalized to 50 µL of 0.1 µg/µL of total peptide, and 30 µL of each sample was transferred to vials (MicroSolv 2 mL vials with AQ Brand deactivated glass low volume inserts) for MS analysis.

### Liquid Chromatography Tandem Mass Spectrometry

Liquid chromatography tandem mass spectrometry (LC-MS/MS) analysis of global proteomics samples was conducted using a Waters nanoAcquity ultra performance liquid chromatography (UPLC) system connected to a Q Exactive Plus Orbitrap mass spectrometer (Thermo Scientific). Samples were loaded into a precolumn (150 μm i.d., 4 cm length, packed in-lab with Jupiter C18 packing material, 300 Å pore size, 5 μm particle size; Phenomenex) using mobile phase A (0.1% formic acid in water). The separation was carried out using a self pack NanoLC column (CoAnn Technologies, 75 µm i.d., 30-cm column) with Waters BEH C18 packing material, 130-Å pore size, 1.7 µm particle size (Waters Corporation). Separations were performed a flow rate of 200 nL/min using a 120 min gradient of 1-75% mobile phase B (ACN + 0.1% formic acid). To prevent carryover, the column was washed with 95-50% mobile phase B for 20 min and equilibrated with 1% mobile phase B for 30 min before the next sample injection.

The mass spectrometer source was set at 2.2 kV, and the ion transfer capillary was heated to 300 °C. The data-dependent acquisition mode was employed to automatically trigger the precursor scan and the MS/MS scans. The MS1 spectra were collected at a scan range of 300-1800 m/z, a resolution of 70,000, an automatic gain control (AGC) target of 3 x 10^6^, and a maximum ion injection time of 20 ms. For MS2, the top 12 most intense precursors were isolated with a window of 1.5 m/z and fragmented by higher-energy collisional dissociation (HCD) with a normalized collision energy at 30%. The Orbitrap was used to collect MS/MS spectra at a resolution of 17,500, a maximum AGC target of 1 x 10^5^, and maximum ion injection time of 50 ms. Each parent ion was fragmented once before being dynamically excluded for 30 s.

### Analysis of Mass Spectrometry Data

Proteomics data have been deposited in MassIVE (Accession code: MSV000096247). **Request for access can be made to the corresponding author.**

MS/MS Automated Selected Ion Chromatogram (MASIC) generator was used to generate selected ion chromatograms (SICs) for all of the parent ions chosen for fragmentation in the LC-MS/MS data[29].

MSGF+ software (v2024.03.26) [30] was then used to perform peptide searches against the *M. tuberculosis* protein database containing 3994 entries (UniProt for Mycobacterium tuberculosis H37Rv, downloaded on 2021-03-07, including the sequence for carbapenem resistance factor [CrfA] Rv2421/Rv2422, downloaded on 2017-10-12) and 16 common contaminant sequences (porcine and bovine trypsin, chymotrypsinogen, human and bovine albumin, and some keratins). Searches were performed with the following parameters: partially and fully tryptic peptides; parent ion tolerance of 20 ppm; methionine oxidation (+15.9949 Da) as a dynamic modification. A spectral probability value of 1.56 x 10^-8^, which was calculated for a 1% false discovery rate (FDR), was used for MSGF+ analysis. This value was determined by calculating reverse hits using the forward+reverse (decoy) database[31] for *M. tuberculosis*.

Global proteomics analyses were performed using the Multi-Omics Analysis Portal and the pmartR package[32]. Potential sample outliers were assessed by a robust Mahalanobis distance (rMd) using the rMd squared values associated with the peptide abundances vector (rMd-PAV), which was applied with all default metrics for proteomics data (correlation, kurtosis, MAD, skewness, and proportion missing)[33]. No sample was identified as an outlier, and all samples were included in analyses. Peptide data were log2 transformed and median normalized. Redundant peptides (those mapping to more than one protein) and peptides found in less than two samples were removed before protein rollup. Peptide ion intensities were rolled up to the protein level using the Rrollup method[34] with median centering. Proteins mapped by less than two peptides were removed. Statistical differences in protein abundance between groups was assessed via analysis of variance (ANOVA) and independence of missingness (IMD, G-test)[35]. In cases when a protein was observed in at least two samples in each group, it was considered significantly different if the mean log2 abundance between groups had a P ≤ 0.05 by ANOVA. In cases when a protein was observed in less than two samples in one group, the number of observed values between groups was considered statistically different if P ≤ 0.05 by IMD.

### Protein Functional Classification and String Analysis

Functional classification of proteins was done using Mycobrowser *M. tuberculosis* H37Rv annotation (Release 5; 2024-07-11) [36]. Data was downloaded from the Mycobrowser website August 2024 (https://mycobrowser.epfl.ch/releases).

Proteomic data was visualized using Cytoscape (ver. 3.10.2)[37] and the StringApp (ver. 2.1.2)[38]. Briefly, the network was created by uploading the UniProt identities of proteins of interest into “STRING: protein query” to retrieve *Mtb* proteins, namely any protein that was up-regulated at least 2-fold in CS conditions (421 proteins). Next, the proteomics data (counts, mean intensity, fold-change, P-value, functional category, etc.) were imported and matched based on the UniProt ID. The ClusterMaker app (ver. 2.3.4) with an MCL partitioning algorithm was used to cluster proteins[39]. Protein nodes were colored based on Mycobrowser functional category. Each node was sized based on the number of CS samples the protein was found in. The final view was restricted to networks that included putative CS biomarkers, which were denoted with a square.

## Results and Discussion

### Global proteomics analysis

We acknowledge that no single model captures all dormancy phenotypes found in patients with TB. In our prior work, we have used the Wayne model, a multi-stress model, and carbon-starvation[27, 40, 41]. For the current work, we used the Grant and Hung model of nutrient starvation[17], which induces an antibiotic-tolerant, nonreplicating state through 5 week incubation in starvation medium (carbon starvation, CS). We cultured six biological replicates of *Mtb* mc^2^6020, a double auxotrophic strain derived from *Mtb* H37Rv[26], to mid-log phase (replicating, Rep) or under CS. After 5 weeks, CS cultures had entered stasis, defined as minimal growth assessed by optical density (**Table S1**).

Global proteomic profiles of whole cell lysates were assessed via liquid chromatography tandem mass spectrometry (LC-MS/MS). We identified 2269 distinct proteins out of the 3994 annotated proteins in *Mtb* (57% proteome coverage) (**Table S2**). A limitation of bottom-up proteomics is a subset of proteins is not detected, particularly for proteins that are low abundance. In *Mtb*, there is over 6 orders of magnitude difference between the most and least abundant proteins [12, 42]. However, the proteome coverage we achieved is on par with other *Mtb* proteomic studies[11, 12, 20, 24]. To minimize false-positives, we only considered proteins identified in at least three of six replicates of either group (Rep or CS), reducing the overall protein count to 2085 (**Figure 1A, Table S2**). Similar protein counts were identified in Rep (1773) and CS (1808) conditions, and most proteins (1496) were identified in both conditions.

**Figure 1.**
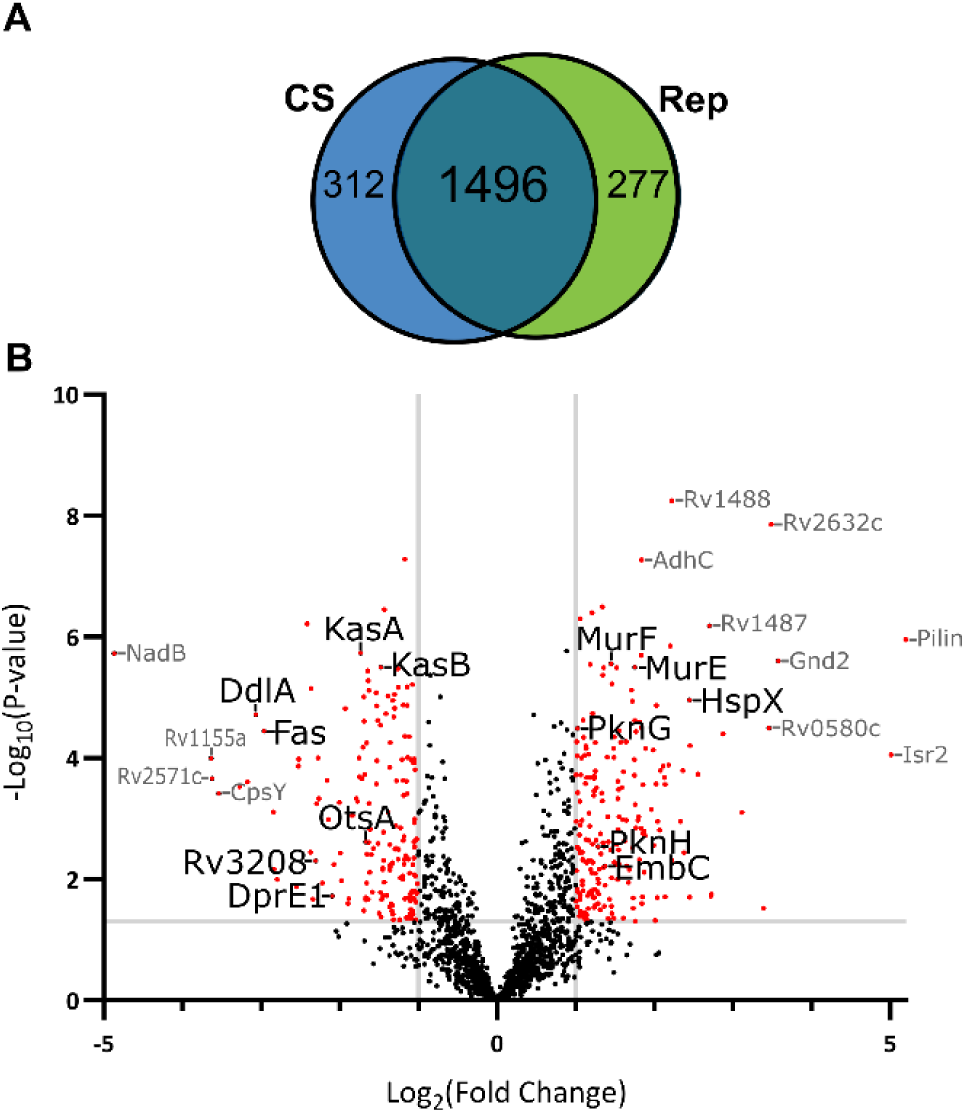
Comparison of proteins identified in CS versus Rep conditions. **A**. Venn diagram of proteins identified in n ≥ 3 samples in CS or Rep conditions. **B**. Volcano plot of proteins found in CS and Rep conditions. Red dots highlight proteins significantly different between groups with P ≤ 0.05 (ANOVA) and a log_2_ fold-change (FC) in CS relative to Rep ≥ 1.0 (upregulated) or ≤ −1.0 (downregulated). Proteins identified in only one group (n ≥ 3 in group A and n ≤ 1 in group B) were not included in the volcano plot due to the inability to calculate a FC. A subset of proteins discussed here are labeled (black font).

We used spectral intensities to estimate abundance and identify proteins that were differentially regulated between growth conditions. We defined differentially-regulated proteins as those found to be significantly different (IMD-ANOVA, P ≤ 0.05) in presence (n ≥ 3 in group A and n ≤ 1 in group B) or abundance (at least two-fold) across groups. There were 415 proteins enriched in CS, with 43.8% not found in Rep lysates (**Figure 1B**, **Table S2**). Conversely, there were 336 proteins enriched in Rep, with 42.8% not found in CS. There were 1142 proteins found to be equally abundant in CS and Rep based on our criteria. Overall, 36% of the quantified *Mtb* proteins were significantly altered in response to carbon starvation, indicating robust and broad reprogramming of cellular functions, organization, and processes.

Identified proteins were assigned to functional categories as defined by Mycobrowser[36] (**Figure 2**, **Table S2**). Differentially abundant proteins were found in all functional categories and, generally, followed patterns of total protein counts (**Figure 2A**). Most of the proteins that were up-regulated in CS were associated with intermediary metabolism and respiration (133; 31%), cell wall and cell wall processes (88; 21%), or conserved hypothetical functions (119; 28%). Proteins that were down-regulated were primarily involved in intermediary metabolism and respiration (76; 23%), cell wall and cell wall processes (43; 13%), and conserved hypotheticals (78, 23%). The representation of functional categories within up- and down-regulated proteins was widely similar, with no category found to undergo large changes in a certain direction (**Figure S1**). The largest categorical shifts with CS were in lipid metabolism (13% down, 5% up) and information pathways (13% down, 4% up).

**Figure 2.**
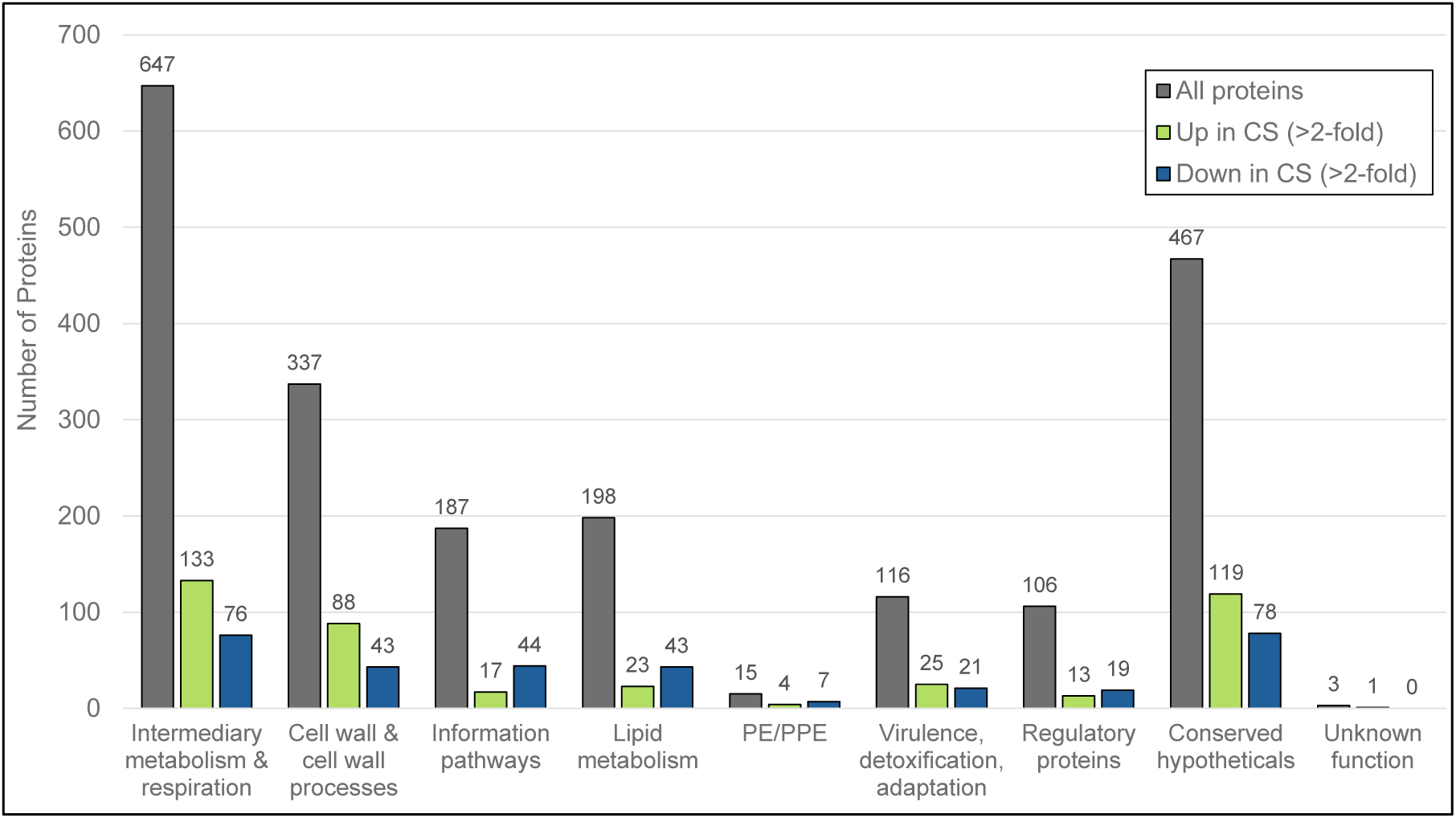
Protein functional classification. Bar chart displaying the distribution of identified proteins across functional categories. The distribution of all proteins found in either Rep or CS conditions is shown in grey (all proteins). The distribution of differentially-regulated proteins in CS relative to Rep are shown in green (up in CS) and blue (down in CS). Categories were defined and assigned based on Mycobrowser [36] *M. tuberculosis* H37Rv (Release 5) annotation.

### Energy Metabolism

Similar to other studies[16, 17, 24], we observed a dramatic reduction in *Mtb* cellular replication with CS (**Table S1**), with little change in optical density (OD_600_) after five weeks of culture. Others have reported significant reductions in cellular respiration rates and intracellular ATP levels with nutrient-starvation models[16, 23]. However, as an obligate aerobe, *Mtb* requires continuous respiration to survive, even in hypoxic and nutrient-starved environments[23, 43]. In accordance with these findings, we observed an overall retention of oxidative phosphorylation enzymes, with many components of the electron transport chain equally abundant or upregulated in CS (**Figure 3A**).

**Figure 3.**
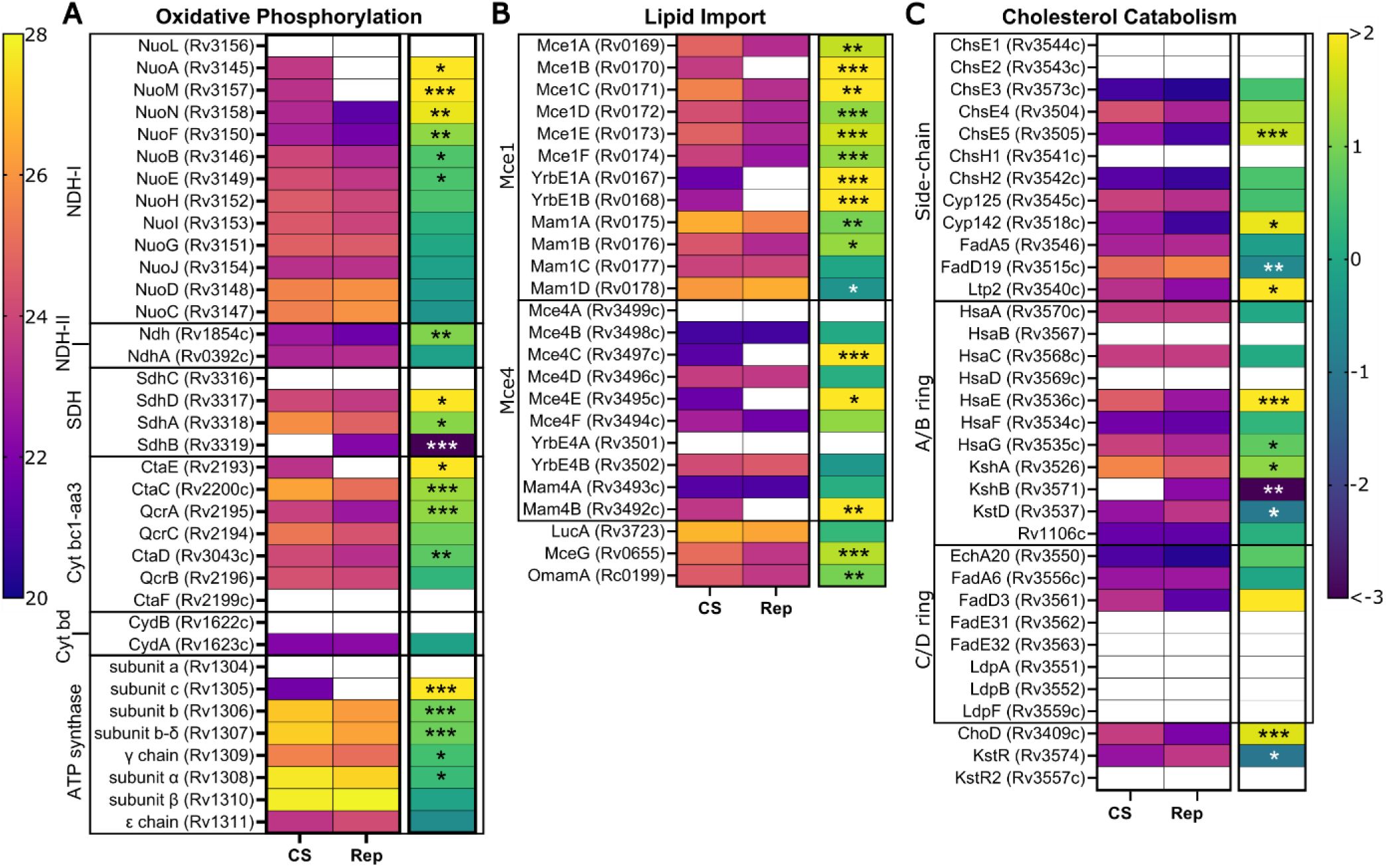
Regulation of energy metabolism in CS versus Rep conditions. Heat maps display group mean intensity values (left map, warm scale) and corresponding FC (right map, cool scale) of proteins in CS versus Rep groups. Proteins are identified by UniProt name and locus ID and are grouped by function: oxidative phosphorylation (**A**), lipid import (**B**), and cholesterol catabolism (**C**). Scale bars are applicable across all corresponding maps. A white box indicates absence of value. The FC of proteins identified in only one group (Group A: n ≥ 3, Group B: n ≤ 1) was set to the respective maximum value. Asterisks denote significance in mean intensity (*: P ≤ 0.05, **: P ≤ 0.01, ***: P ≤ 0.001).

In contrast to findings in hypoxia-induced models of dormancy[44], oxidative phosphorylation appears to progress through the aerobic chain with many components of the cytochrome *bc1*-*aa3* complex being upregulated in CS. No significant alteration to the terminal oxygen acceptor of the anaerobic chain, cytochrome *bd* oxidase (cyt *bd*), was observed. While increased reliance on cyt *bd* has been associated with *Mtb* adaptation to hostile environments, this reliance appears to be related to the oxygen-deficient and oxidative stress conditions within the host[45, 46]. We did not observe this same pattern in our nutrient-starvation model, highlighting a key difference between hypoxia and nutrient-starvation models.

Key components of NADH dehydrogenase type I (NDH-I) and ATP synthase complexes were more abundant in CS. Of particular interest is the increased abundance of ATP synthase subunit c (Rv1305), the direct target of bedaquiline[47, 48]. This finding aligns with previous studies demonstrating that dormant bacilli have increased susceptibility to bedaquiline and adds further understanding to the efficacy of this drug against TB[49].

### Lipid uptake and catabolism

During infections, the primary carbon and energy source for *Mtb* is the co-utilization of host cholesterol and simple carbon fatty acids[50]. An interesting finding in our CS model is the significant upregulation of the Mce1 and Mce4 complexes, which, respectively, import fatty acids and cholesterol into the *Mtb* cell (**Figure 3B**). Both of these ABC transporter complexes are comprised of six Mce proteins (e.g., Mce1A-F) and two YrbE proteins (e.g., YrbE1A/B), and are associated with different numbers of Mam accessory proteins[51, 52]. Every protein of the core Mce1 transporter structure is upregulated in CS. Several of the Mam1 proteins are also more abundant in CS, as well as the ATPase common to all Mce transporters, MceG (Rv0655). The cholesterol transporter Mce4 also appears more abundant in CS, with Mce4C (Rv3497c) and Mce4E (Rv3495c) only found in CS and Mce4D (Rv3496c) and Mce4F (Rv3494c) found more consistently across the CS group than Rep (**Table S2**). While the implications of Mce1 complex upregulation are still not fully understood, it is known that cholesterol uptake by the Mce4 complex is crucial to survival of the pathogen in nutrient-starved host environments [53, 54]. Our data indicate that *Mtb* compensates for CS by enhancing lipid uptake.

*Mtb*’s requirement for cholesterol during CS suggested to us that cholesterol catabolism would be upregulated in CS. *Mtb* uses over 30 enzymes to catabolize cholesterol, as reviewed[55, 56]. Twelve enzymes degrade the side-chain (Cyp125/142, ChsE1/2/3/4/5, ChsH1/H2, FadA5, FadD19, and Lpt2). Of the nine found in our study, most were up-regulated in CS (**Figure 3C**). Twelve enzymes degrade the A and B rings of cholesterol: ChOX (Rv1106c), KtsD (Rv3537), KshA/B (Rv3526/Rv3571), and HsaA/B/C/D/E/F/G/H. We identified nine of these and only two (KstD and KshB) were downregulated in CS. Three were upregulated in CS (KshA, HsaE, and HsaG). C/D ring catabolism involves at least 8 enzymes; most were not found in our data. Catabolism of the A/B ring is negatively regulated by a transcriptional repressor, KstR1 (Rv3574)[57], which was > 2-fold down-regulated in CS. Within our data, we identified 53 proteins that are putatively regulated by KstR1. Nineteen were up-regulated in CS, including Mce4C, HsaE/G, KshA, and Ltp2. Two KstR1-regulated proteins, OtsB1 (Rv2006) and EchA19 (Rv3516), were identified only in CS, and we suggest that these two might serve as CS biomarkers (see below). Overall, our data supports the important role that cholesterol degradation plays in *Mtb* survival in nutrient-limited environments.

*Mtb*’s genome encodes over 300 proteins involved in lipid metabolism[36, 58]. There are many lipolytic enzymes, including esterases, lipases, cutinases, and phospholipases[59]. Prior work in our group[27] and by others[60, 61] demonstrated that lipase activity is regulated in hypoxia. In the current work, we identified 14 lipases (Lip family members), 1 cutinase (Culp6), and 13 additional esterases (**Table S3**). Many of these enzymes were differentially abundant in CS (7 up, 13 down), further emphasizing the importance of lipid metabolism regulation in stasis.

### Regulation of serine/threonine protein kinases

*Mtb* has eleven serine/threonine protein kinases (STPKs). Recent work by Frando and coworkers demonstrated the importance of the *Mtb* STPKs in widespread *o*-phosphorylation of the proteome[62]. Specifically, they showed that over 70% of mycobacterial proteins are *o*-phosphorylated and that 30% of *Mtb* gene expression was changed by perturbing STPK levels. We did not examine phosphorylation of proteins here. However, we detected nine of the eleven STPKs in our analysis (**Table S3**). We found that the two essential STPKs, PknA (Rv0015c) and PknB (Rv0014c), were identified in all samples. PknE (Rv1743) and PknJ (Rv2088) were found only in Rep samples, indicating strong down-regulation in CS.

Notably, we found PknG (Rv0410c) and PknH (Rv1266c) were significantly up-regulated in CS. Both of these STPKs are linked to metabolic adaptation and long-term survival in dormancy[63]. PknG facilitates survival in hypoxia[64] and under oxidative stress[65]. PknG responds to nutrient stress[66], through TCA cycle regulation via phosphorylation of glycogen accumulation regulator A (GarA; Rv1827)[67]. GarA was up-regulated 2-fold in CS. Furthermore, PknG activity is linked to trafficking and survival in host macrophages[68]. PknH was up-regulated 2.4-fold in CS. This STPK restricts *Mtb* growth *in vivo*[69]. PknH phosphorylates DosR (Rv3133c) and up-regulates the DosR regulon in response to nitric oxide[70]. Overall, the observed up-regulation of PknG and PknH aligns with their previously reported roles in mediating survival of dormant *Mtb*.

### Cell wall biosynthesis is largely retained in CS

*Mtb* has an unusual cell wall that provides challenges and opportunities for treatment of TB[71–73]. The mycobacterial cell wall is composed of three layers: peptidoglycan (PG), arabinogalactan (AG), and mycolic acids (MA). Other components include glycophospholipids, sulfolipids, trehalose, and phosphatidyl-myo-inositol mannosides. Dormant *Mtb* have altered cell envelopes, with loss of acid fast staining[74]. The cell wall has reduced MA[13], trehalose monomycolate, trehalose dimycolate, and phosphatidylinositol mannosides (PIMs). On the other hand, the cell wall has increased lipomannans and lipoarabinomannans in dormancy[8]. *Mtb* PG has both classical 4→3 crosslinks and non-classical 3→3 crosslinks. It is not yet clear if *Mtb* PG crosslinking is altered in dormancy, although the PG in replicating cells likely has fewer 3→3 crosslinks than the PG in stationary phase cells (80%)[21] or hypoxic cells (∼70%)[75]. For these reasons, we assessed the regulation of the various enzymes involved in the biosynthesis of the cell wall in response to CS.

We examined the enzymes involved in PG biosynthesis and found that many were unchanged between CS and Rep except for a few key players (**Figure 4A**). PG is comprised of a repeating glycan backbone of N-acetylglucosamine (GlcNAc) and N-acetylmuramic acid (MurNAc) interspersed with crosslinked peptide stems. PG biosynthesis starts in the *Mtb* cytoplasm with the synthesis of UDP-GlcNAc conjugated to the peptide stem (D-Ala—D-iso-Gln—*meso*-DAP—D-ala—D-ala). The glycan portion is synthesized by GlmS (Rv3436), GlmM (Rv3441c), GlmU (Rv1018c), MurA (Rv1315c), and MurB (Rv0482c); all were identified in both CS and Rep samples. The peptide stem is formed by MurC/D/E/F. Both MurE (Rv2158c) and MurF (Rv2157c) were up-regulated >2-fold in CS. The D-ala—D-ala ligase, DdlA (Rv2981c), was down-regulated (>8-fold) in CS. There are modifications to the nascent PG that are mediated by membrane proteins, but many of these were not identified in our samples (e.g., NamH, MurX/MraY, MurJ, MurG). AsnB (Rv2201) was down-regulated and the putative flippase FtsW (Rv2154c) was absent in our CS samples. CwlM (Rv3915) regulates MurA activity[76]; it was only found in Rep samples.

**Figure 4.**
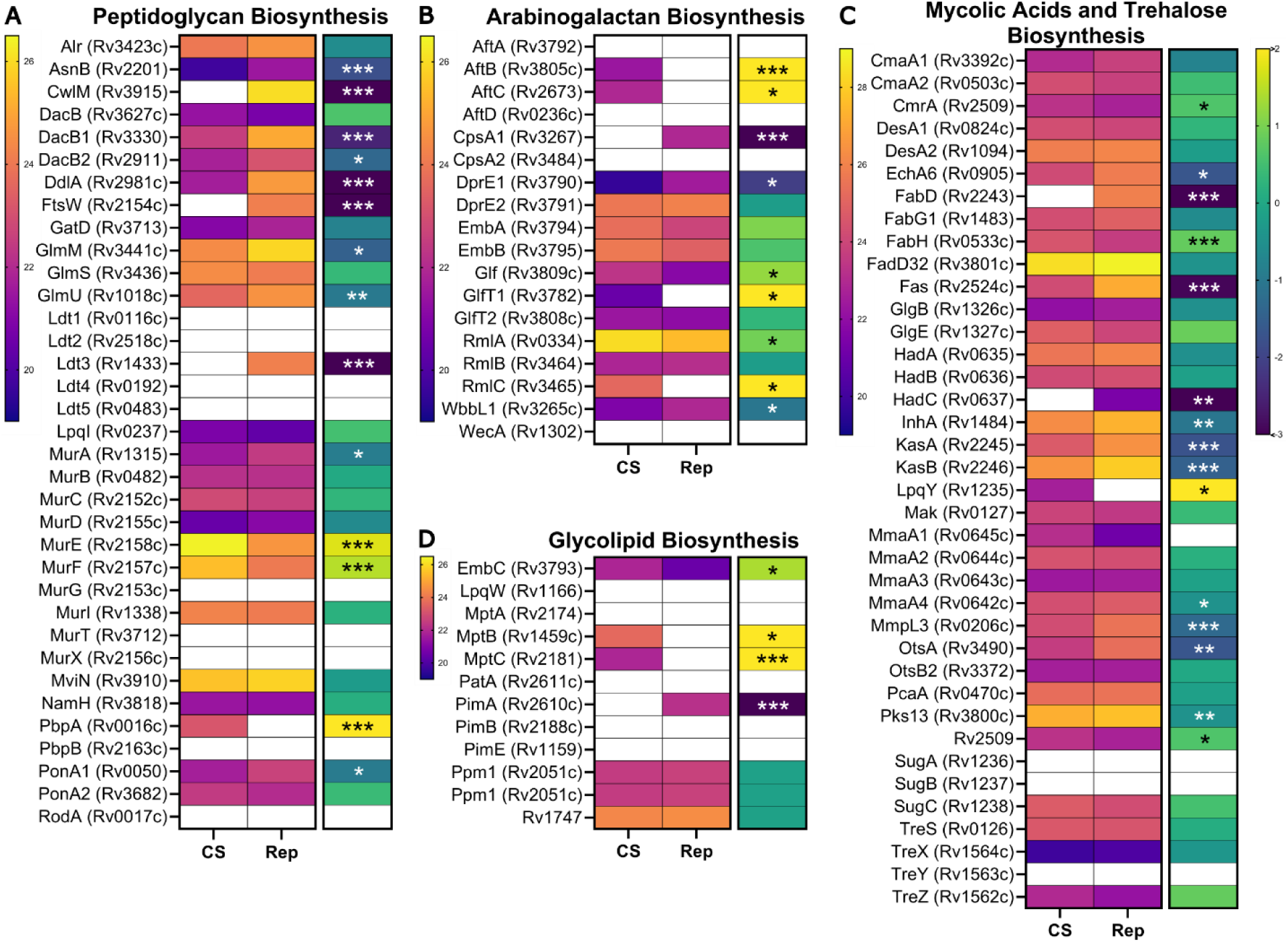
Changes in cell wall biosynthesis in response to CS. Proteins are identified by Uniprot name (locus ID) and grouped by function: PG biosynthesis (**A**), AG biosynthesis (**B**), MA and trehalose biosynthesis (**C**), and glycolipid biosynthesis (**D**). Heat maps display group mean intensity values (left map, warm scale) and corresponding FC (right map, cool scale) of proteins in CS versus Rep groups. Fold change scale bar (cool scale) is applicable across all FC maps. A white box indicates absence of value. The FC of proteins identified in only one group (Group A: n ≥ 3, Group B: n ≤ 1) was set to the respective maximum value. Asterisks denote significance in mean intensity (*: P ≤ 0.05, **: P ≤ 0.01, ***: P ≤ 0.001).

The remainder of PG biosynthesis occurs in the periplasm, where the activity of various penicillin binding proteins (e.g., PonA1, PonA2, PBPA/B), L,D-transpeptidases (e.g., Ldt1-5), and carboxypeptidases (e.g., DacB1/2) produce the final crosslinked PG structure[77]. The abundance of most identified penicillin binding proteins was unchanged. Interestingly, DacB1 and DacB2 were both significantly down-regulated in CS. Their function is to remove a D-Ala from the peptide stem, a reaction that precedes Ldt (3→3) crosslinking[75]. We speculate that down-regulation of carboxypeptidases in CS would result in fewer tetrapeptide stems available for Ldt crosslinking. Only one Ldt, Ldt3 (Rv1433), was detected in our samples (Rep), presumably because the abundance of other Ldts was below the MS limit-of-detection. Although Betts found that Ldt1 (Rv0116c) was transcriptionally up-regulated (17-fold) in dormancy[16], our work and several other proteomics studies have been unable to detect Ldt1 at the protein level[24, 28]. Prior studies characterized PG crosslinking in replicating and stationary-phase *Mtb*, finding that 70-80% of the PG is 3→3 crosslinked in an Ldt-dependent manner[21, 75, 78]. To our knowledge, the composition of PG crosslinks in nutrient-starved *Mtb* has not been studied. Our data suggest that further investigation is warranted to characterize PG crosslinks in CS-induced dormancy.

Arabinogalactan (AG) is a polymer of branched arabinose chains attached to a “trunk” of galactose[72]. AG is covalently attached to both PG and MA. Biosynthesis of AG starts in the cytosol with several enzymes forming the galactan chain: WecA (Rv1302), WbbL (Rv3265c), and GlfT1/T2 (Rv3782/Rv3808c). We found that WbbL was down-regulated in CS, while GlfT1 was only observed in CS, indicating strong upregulation (**Figure 4B**). Arabinose is added to the galactose trunks in the periplasm by arabinosyltransferases: AftA (Rv3792), AftB (Rv3805c), AftC (Rv2673), AftD (Rv0236c), and EmbA/B (Rv3794/Rv3795). AftA and AftD were not found in our samples, while AftB and AftC were only found in CS. EmbA was up-regulated (>2-fold) in CS. Two other relevant AG biosynthetic enzymes are DprE1/E2 (Rv3790/Rv3791). DprE1 was down-regulated in CS (>4-fold) while DprE2, a putative target of pretomanid[79], was unchanged. These findings suggest that AG biosynthesis is altered under CS conditions.

The MA layer is comprised of long-chain fatty acids (e.g., trehalose mono- and di-mycolate) that form a protective hydrophobic barrier. Many of the enzymes associated with de novo MA biosynthesis were down-regulated in our CS samples (**Figure 4C**). The key fatty acid synthesis enzyme FAS-I (Rv2524c) was strongly down-regulated in CS (>7-fold), as was the *fas* transcriptional regulator Rv3208 (∼5-fold)[80]. Newly synthesized fatty acids are extended by the FAS-II complex of enzymes. Several FAS-II enzymes were down-regulated, including KasA/B (∼3-fold; Rv2245/Rv2246) and InhA (2-fold; Rv1484). Prior work demonstrated that loss of KasB prevents acid-fast staining[81]. HadC (Rv0637) was only identified in Rep samples; HadA/B (Rv0635/Rv0636) were unchanged. Trehalose monomycolate, a MA, is transported to the periplasm via MmpL3 (Rv0206c); this essential protein was down-regulated in CS (>2-fold). Proteins involved in mycolic acid synthesis have also been found to be downregulated at the transcript and protein level under hypoxic conditions[12, 82]. Overall, these patterns across multiple *in vitro* models suggest it is beneficial to dormant *Mtb* to suppress energy-demanding MA biosynthesis[53, 83]. Attachment of MA to AG is mediated by the antigen 85 complex (Rv3804, Rv1886c, Rv0129c), which was not detected in any of our samples.

We observed several CS-related changes in enzymes that synthesize the glycolipids and lipoglycans of the inner and outer membrane-phosphatidylinositol mannosides (PIMs), lipomannans (LMs), and lipoarabinomannans (LAMs) (**Figure 4D**). It has been proposed that PIMs are less abundant in stasis while LMs and LAMs are upregulated[8]. We saw concordance with this theory in our data. The α-mannopyranosyl transferase that initiates PIM biosynthesis, PimA (Rv2610c), was only found in Rep samples, indicating strong downregulation with CS. We also found hints that LM and LAM biosynthesis were upregulated. The mannosyltransferases, MptB (Rv1459c) and MptC (Rv2181), were both only found in CS samples, indicating strong upregulation. These enzymes are involved in the processing of PIMs to LMs, while the arabinotransferase EmbC (Rv3793) is responsible for modifying mature LM with arabinose to form LAMs[72, 84]. We found EmbC was also significantly more abundant in CS (5.5-fold).

### Drug targets in CS and Rep *Mtb*

Drug treatment for TB is long, usually lasting six months or more[1, 5]. Monotherapy is not used for *Mtb* due to low efficacy and the high probability of selecting drug-resistant mutants. The most common treatment regimen for drug-susceptible TB includes isoniazid, rifampicin, pyrazinamide, and ethambutol. For years, the treatment of drug-resistant disease varied patient to patient, with generally poor outcomes (15-40% patient mortality globally)[1]. Treatment outcomes for drug-resistant TB improved in 2022, with approval of the combination therapy of bedaquiline, pretomanid, and linezolid (BPaL)[85, 86]. Other drugs used to treat TB include fluoroquinolines, rifapentine, cycloserine, streptomycin, kanamycin, and (rarely) β-lactams[87]. Currently, standard treatment for LTBI is rifampicin, rifampicin plus isoniazid, or rifapentine plus isoniazid[92].

Nearly all (99.9%) of *Mtb* are killed within the first two weeks of treatment[88]. The remaining months of treatment are implemented to kill non-replicating *Mtb* that are particularly challenging to eradicate and the focus of our study. Many studies have found that antibiotics have decreased potency against dormant *Mtb*[16, 17, 22, 89, 90]. This phenomenon is termed phenotypic drug resistance and is a hallmark of *in vitro* dormancy models[6]. A 2017 review summarized findings from the literature and found that rifampicin, rifapentine, metronidazole, bedaquiline, pretomanid, and fluoroquinolines are the most active drugs against non-replicating *Mtb*[91]. Studies have found that nutrient-starved *Mtb* is more drug tolerant than hypoxic *Mtb*[16, 17, 23, 90]. For example, rifampicin is ∼50-fold less potent against nutrient-starved *Mtb* compared to hypoxic *Mtb*[23].

We analyzed our proteomic data for the relative levels of drug targets in CS and Rep *Mtb* samples (**Figure 5**). The *Mtb* cell wall is a target of many antibiotics. Synthesis of mycolic acids is inhibited by isoniazid and ethionamide, which both target an enoyl-[acyl-carrier-protein] reductase (InhA; Rv1484)[93]. This enzyme was found in both CS and Rep samples, but was down-regulated in CS (2-fold). The levels of the enzymes that activate isoniazid and ethionamide, KatG (Rv1908c) and EthA (Rv3854c) respectively, were unchanged. A trehalose monomycolate exporter, MmpL3 (Rv0206c), is the target of SQ109[94], a drug that is still under evaluation for treatment of TB. It was down-regulated >2-fold in CS.

**Figure 5.**
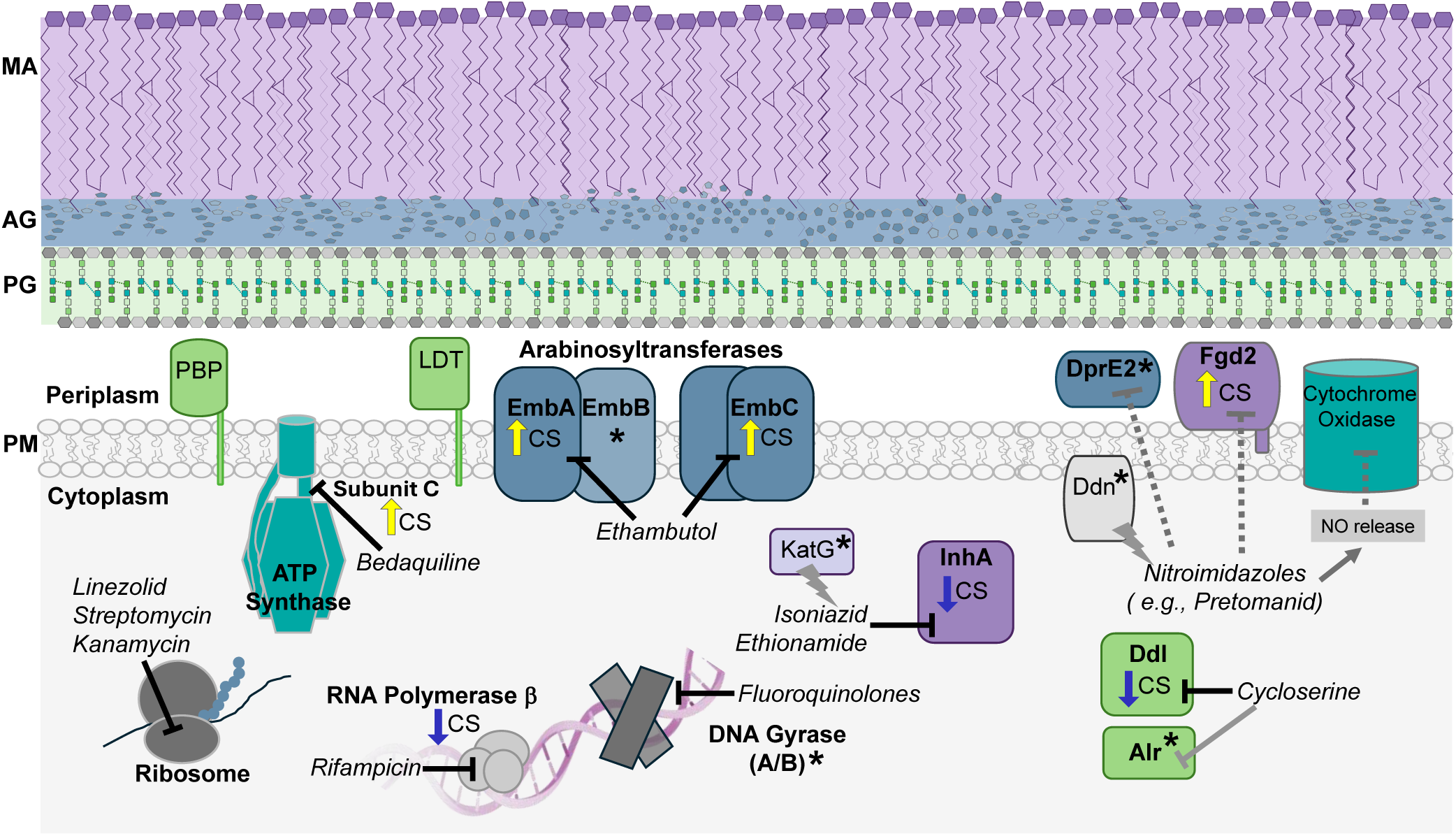
The targets of clinically approved TB drugs are present in *Mtb* under CS and Rep conditions. Drug targets involved in cell wall biosynthesis are color-coded to indicate their downstream target: mycolic acid (MA; purple), arabinogalactan (AG; blue), peptidoglycan (PG; light green). Proteins involved in transcription and translation are shown in gray. Other relevant drug targets are shown in teal. Altered protein levels in CS are indicated by arrows: yellow (up-regulated) and dark blue (down-regulated). An asterisk (*) indicates no significant change in protein levels between CS and Rep. Nitroimidazoles have a complex mechanism and target more than one pathway, as indicated.

Nitroimidazoles (e.g., pretomanid and delamanid) have a complex mechanism of action[95]. Pretomanid is effective on replicating and non-replicating *Mtb*. In hypoxia, pretomanid is activated by deazaflavin-dependent nitroreductase (Ddn; Rv3547) and produces reactive nitrogen species (e.g., nitric oxide, NO), which inhibits cytochrome oxidase[95]. Pretomanid may also inhibit a F420-dependent hydroxymycolic acid dehydrogenase (Fgd2; Rv0132c)[96]. This enzyme is involved in MA biosynthesis and was only identified in our CS samples. We propose this enzyme as a CS biomarker (see below). Pretomanid may inhibit DprE2 (Rv3791)[79], an essential enzyme that forms a precursor for AG. Protein levels of Ddn and DprE2 were unchanged in CS versus Rep conditions, suggesting that these targets are drug-susceptible in nutrient-starved *Mtb*.

The arabinosyltransferases EmbA (Rv3794), EmbB (Rv3795), and EmbC (Rv3793) polymerize arabinose during AG biosynthesis and are inhibited by ethambutol, a front-line drug. The A and C subunits were both up-regulated >2-fold in CS, signifying that AG biosynthesis is important during CS. Yet prior studies found that ethambutol was less effective in nutrient-starved and hypoxic *Mtb*, which suggests that access to these transferases is reduced[9, 89, 90]. Cycloserine targets PG biosynthesis through inhibition of the D-alanine—D-alanine ligase (Ddl; Rv2981c) and, to a lesser extent, alanine racemase (Alr; Rv3423c)[87]. Ddl was highly down-regulated in CS (>8-fold). This is consistent with the finding by Xie et al. that cycloserine was not effective in killing nutrient-starved *Mtb*[90]. PG biosynthesis is disrupted by β-lactams, although this therapeutic class is only rarely used to treat TB[97, 98]. Targets of carbapenems, a sub-class of β-lactams, include PBPs and Ldts. Regulation of these enzymes in CS is summarized in **Figure 4A**.

Some drugs target the pathogen’s transcription and translation[87]. Fluoroquinolones (e.g., moxifloxacin) target DNA gyrase A and B (Rv0005 and Rv0006)[99]; both were not significantly changed between CS and Rep samples. Rifampin inhibits the RNA polymerase β-subunit (Rv0667). This protein was down-regulated in CS (2-fold), which perhaps explains *Mtb*’s rifampin resistance in CS conditions[16]. Streptomycin, kanamycin, and linezolid, an oxazolidinone, all target ribosomal RNA[87], which was not quantified in our study.

Bedaquiline is a diarylquinoline that targets ATP synthase, disrupting energy metabolism[47–49]. More specifically, it binds to the C-subunit (Rv1305) of the ATP synthase F_0_ domain. A surprise in our study is that subunit C was only detected in our CS samples. There are 8 subunits of the *Mtb* ATP synthase and subunit C is the only one absent in Rep samples. We hypothesize that subunit C was present but undetectable by MS in Rep samples since ATP production and oxidative phosphorylation are required during normal replication.

Lastly, pyrazinamide is a pro-drug activated by the pyrazinamidase PncA (Rv2043c)[100]. We did not detect this enzyme in our samples. Additionally, there are reports that this drug acts on pantothenate biosynthesis[101]. We used an auxotrophic strain with disruption in two genes that make pantothenate (PanC and PanD)[26]. Therefore, we could not assess regulation of these targets in our study.

To summarize, we found that the targets of many drugs used to treat TB are present in both Rep and CS samples. Before new drugs and drug regimens are approved, it is critical to assess how effective treatments are against dormant *Mtb*. Our analysis provides an initial evaluation of the link between protein abundance in dormancy and drug susceptibility. We view this information as supplemental to direct measurements of drug potency in cells and animal models, since many factors influence efficacy besides target abundance, including efflux pumps and cell wall permeability[89].

### Comparison with prior CS -omics studies

To our knowledge, no previous study has thoroughly characterized whole-cell proteomic changes in *Mtb* in response to extended CS. Our findings presented here fill this gap and provide insight relevant to dormant *Mtb*. Two prior studies have conducted proteomics analysis of *Mtb* under CS[16, 24], albeit with major limitations. Betts and coworkers investigated proteomic changes in *Mtb* in response to short duration CS (5 days); however, they identified only seven proteins as differentially regulated[16].

Matching their findings, we found Tig (Rv2462c) and GrpE (Rv0351) to be down-regulated and Rv2557 and HspX (Rv2031c) to be up-regulated in CS. We saw discordant results for two other proteins (Rv1860 and Rv1980c). The remainder of their ground-breaking work focused on transcriptional changes after four to 96 hours of CS; we will not discuss those results here because of the differences in study design. There have been many advances in MS-based proteomics since the early 2000’s that have greatly improved the power of peptide detection. Still, more recent investigations into nutrient-starved *Mtb* have limitations. The Grant et al. CS study[17] did not include proteomic analysis and the most recent CS study that did was published by Albrethsen et al. in 2013[24]. Albrethsen induced dormancy using the Betts nutrient deprivation model (i.e., standing cultures grown 6 w in PBS). They isolated secreted proteins from culture filtrates for identification using LC-MS/MS analysis but did not analyze whole cell lysates, a limitation of their study.

Despite significant differences in study design, we compared our CS proteomic profile to that from the Albrethsen study [24] (**Figure 6**). As expected, when analyzing cellular proteins compared to secreted proteins, we achieved higher proteome coverage (52% versus 33%, respectively). There were 1001 proteins identified in both of our studies (**Figure 6A**). In Albrethsen’s work, of the 1305 proteins that were identified in at least two samples from either CS or Rep (N = 3), there were 343 up-regulated (>2-fold; P < 0.05) and 288 down-regulated (>2-fold; P < 0.05) in CS. By comparison, we identified 415 up-regulated (>2-fold; P <0.05) and 336 down-regulated (>2-fold; P < 0.05) proteins. These results are illustrated in **Figure 6B**. There were 59 up-regulated and 35 down-regulated proteins identified in both studies, which we consider high confidence CS-responsive proteins (Table S4). There were 51 proteins that showed opposite regulation; we term these discordant. Given the major study differences, it is perhaps surprising that >9% of proteins identified across both studies were concordant. We conclude that the lack of proteomics analyses of the CS model over the last decade highlights the current study as a significant resource.

**Figure 6.**
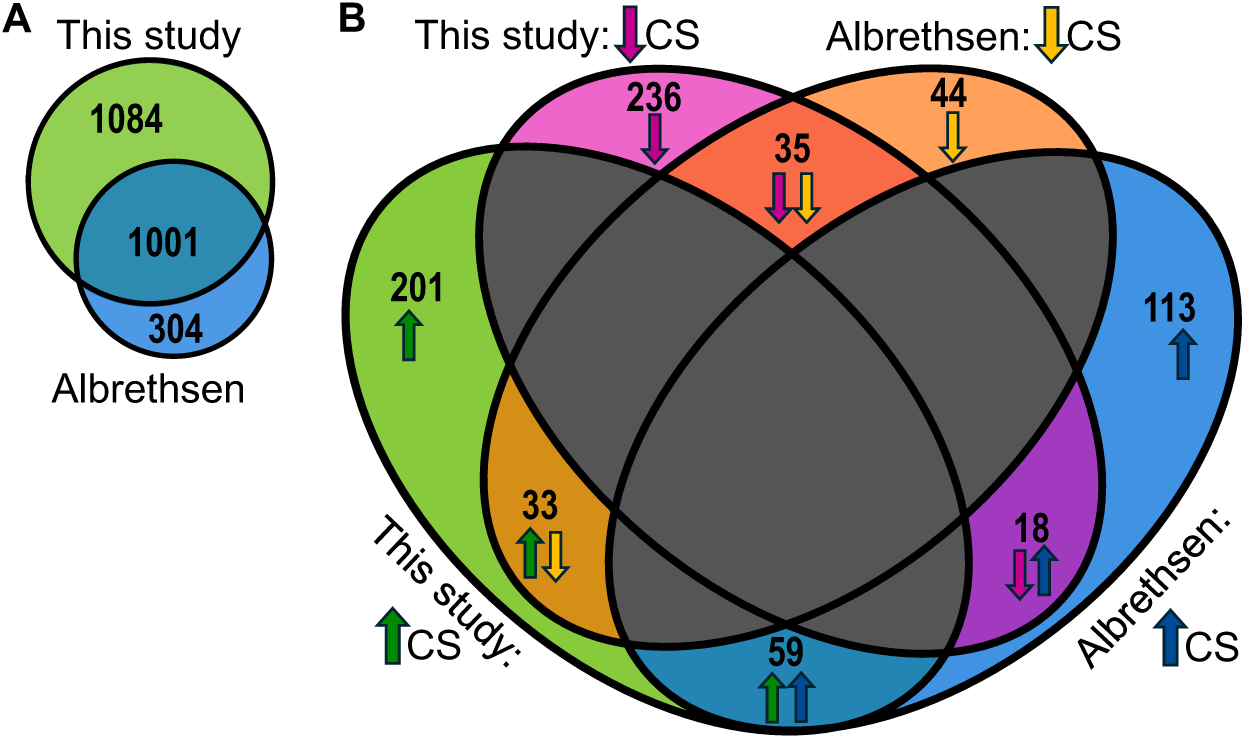
Comparative analysis of proteins associated with nutrient starvation. **A**. Total shared and unique *Mtb* proteins identified in at least three replicates (our study) or 2 replicates (Albrethsen et al. 2013[24]) of either CS or Rep conditions. **B**. Comparison of differentially expressed proteins between the two studies. There were 59 proteins that were up-regulated (FC > 2) and 35 that were down-regulated (FC > 2) in CS in both studies; 51 proteins were discordant between the studies.

### Biomarkers for CS dormancy

Biomarkers for dormancy, especially CS, are variable and poorly defined[7]. For example, a putative biomarker for dormancy is HspX (Rv2031c). Also named alpha crystalline, this protein is a chaperone induced during CS[16] and hypoxia[11, 102]. Betts described the up-regulation of the select isoforms of HspX under CS[16]. However, the use of HspX as a putative dormancy biomarker should be done with extreme caution. It is highly expressed in Rep cultures and is the second most abundant protein in the *Mtb* proteome[42]. HspX was >5-fold up-regulated in our CS samples but was also present in all Rep samples. Lastly, Albrethsen et al. did not detect a significant change in HspX in their CS study[24]. Unless a dormancy-associated isoform of HspX is characterized and validated, we would not recommend using HspX as a dormancy biomarker.

We sought to create a list of putative CS-associated biomarkers composed of proteins highly upregulated in both Albrethsen’s[24] and our data. For our data, we selected proteins found only in CS and absent in Rep, making them high confidence CS markers (182 proteins). We compared this list to proteins that were at least 10-fold up-regulated in Albrethsen’s CS model to identify common proteins[24]. From this analysis, we propose that Fgd2 (Rv0132c), Rv0571c, Rv1019, Gmk (Rv1389), CysG (Rv2847c), FadD13 (Rv3089), EchA19 (Rv3516), and Rv3618 are potential CS biomarkers.

The DosR regulon plays a key role in non-replicating *Mtb* induced by hypoxia[102, 103]. Although we identified 24 DosR-regulated proteins in our study, only nine were up-regulated (>2-fold) in CS. Among those, four DosR-regulated proteins—Rv0571c, Rv2004c, OtsB1 (Rv2006), and Rip3 (Rv2625c)—were present in our CS samples and absent in Rep samples. We propose that these could serve as CS biomarkers, although only one (Rv0571c) was identified as significantly up-regulated in Albrethsen’s study[24]. None of the components of the regulon’s activation system, DosS (Rv3132c), DosT (Rv2027c), and DosR (Rv3133c), were significantly changed between our CS and Rep samples. The stringency response protein, RelA (Rv2583c)[104], was slightly down-regulated (1.8-fold; p = 0.006) in CS, making it a poor biomarker for CS.

We performed network analysis in Cytoscape[38] of all proteins that were up-regulated (P ≤0.05, FC > 2) in CS samples (**Figure 7**). We color-coded proteins based on functional category and limited the display to clusters containing the eleven proteins that we propose to be CS biomarkers: Fgd2, Rv0571c, Rv1019, Gmk, CysG, FadD13, EchA19, Rv3618, Rv2004c, OtsB1, and Rip3. Four of the proteins are implicated in intermediary metabolism and respiration: Fgd2, Gmk, CysG, and Rv3618. Among these, Fgd2 is intriguing because it produces keto-mycolic acids[96], a component of the cell wall. It is a F420-dependent hydroxy-mycolic acid dehydrogenase and a putative target of the drug pretomanid[96]. The functional significance of up-regulation of Fgd2 is unclear, but it could be involved in remodeling mycolic acids in dormancy. A recent study found that most mycolic acids are less abundant in hypoxia-induced dormancy, although there was increased abundance of a C_77_ keto-mycolic acid[13].

**Figure 7.**
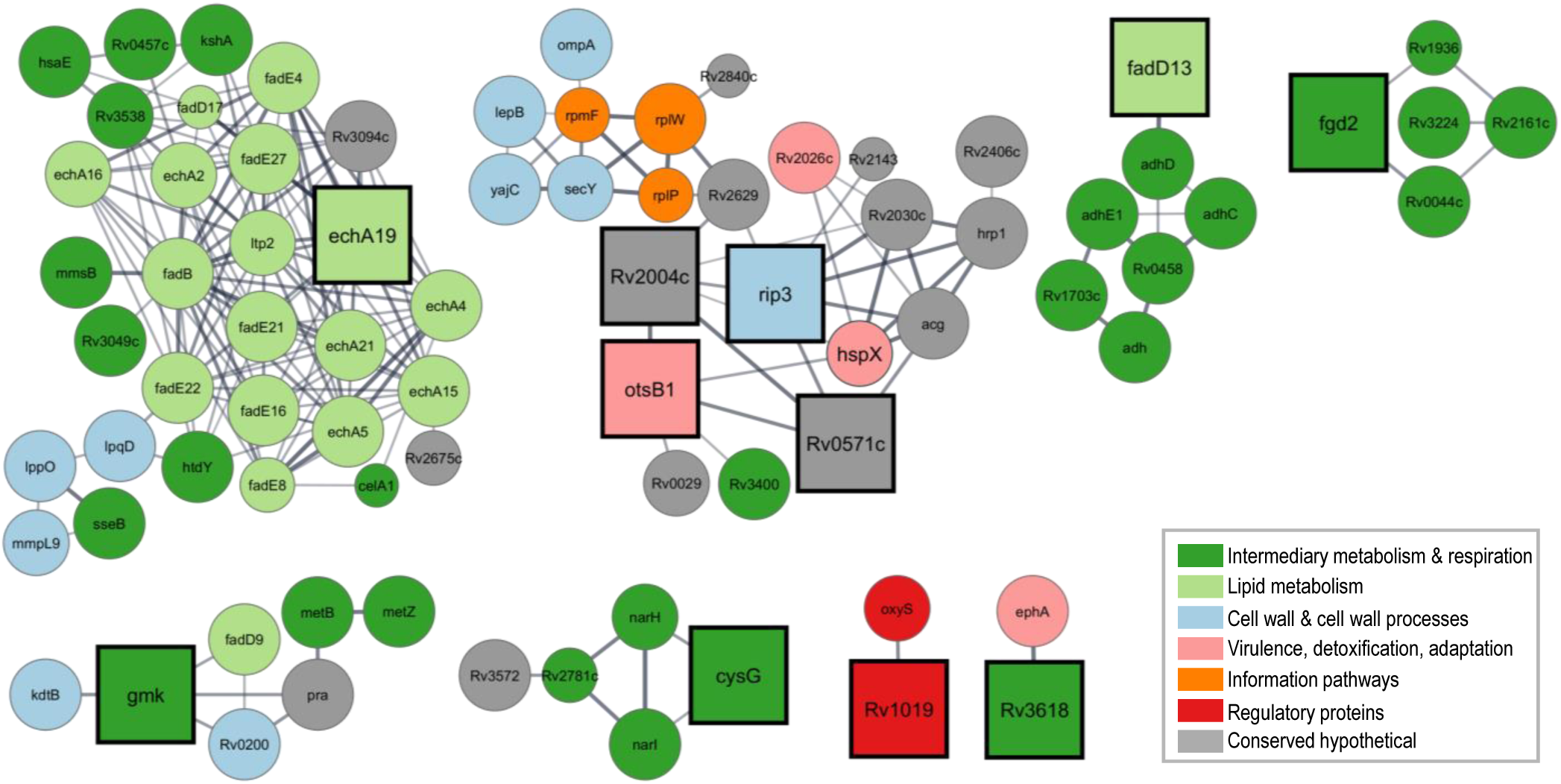
Network and functional analysis of putative CS-associated biomarkers in relationship to other proteins enriched (P ≤ 0.05, FC > 2) in CS. Protein interactions were mapped in Cytoscape using the StringApp and ClusterMaker plug-ins. CS-associated biomarkers are shown as squares. Node size is based on the number of hits in CS (n = 3-6) and node color is coded by Mycobrowser functional category.

FadD13 and EchA19 are linked to lipid metabolism. FadD13 is a long-chain fatty acyl co-A synthetase implicated in the maintenance of the mycolic acids, including under acidic conditions[105, 106]. EchA19 is an enoyl-CoA hydratase involved in cholesterol degradation[107]. OtsB1(Rv2006) is a part of the DosR regulon and may participate in trehalose biosynthesis. It is a putative trehalose-6-phosphate phosphatase with homology to the essential enzyme OtsB2 (Rv3372)[108]; OtsB1 has yet to be biochemically characterized. A 2004 paper claiming that OtsB1 lacked phosphatase activity provided no data to support that conclusion[109]. As noted above, OtsB1 and EchA19 are both KstR1-regulated proteins[57].

We note that several of the proposed CS biomarkers are linked to the cell wall components: Fgd2, FadD13, and OtsB1. It is tempting to speculate that these enzymes facilitate the remodeling of the cell wall for survival in dormancy. Further studies are warranted. As far as we could determine, there are no antibodies available for these proposed CS biomarkers. Future work should focus on generating such reagents for the community to validate these putative CS biomarkers.

## Conclusions

There are several *in vitro* models that induce a non-replicating state in *Mtb* that mimics the bacterial physiology observed during latent TB infections. In the current study, we used carbon-starvation to model *Mtb* dormancy. We report the first in-depth analysis of *Mtb* whole-cell proteomic changes in response to CS. We suggest a set of biomarkers for the CS state that could be used to assess nutrient starvation phenotypes in future studies.

Many of our findings align with previous reports of hypoxia-induced changes, strengthening our understanding of the *Mtb* non-replicative state. In both models, the kinases PknG and PknH play a central role in responding to external stressors. Additionally, we confirmed that nutrient starvation, like hypoxia, influences cell wall biosynthesis with measurable down-regulation of the MA biosynthetic machinery. This result is consistent with effects on *Mtb*’s cell wall and cell wall staining observed in patient samples[110]. However, we also observed key differences in protein regulation between CS and previous hypoxia-related findings. For example, energy metabolism appears to proceed through the aerobic chain in CS. We also found evidence that lipid uptake and catabolism is upregulated in CS, which has not been described for hypoxia models. Furthermore, the DosR regulon appears much less central to CS-induced reprogramming than in hypoxia.

Considering the difficulty of modeling the complex and multifaceted environment that dormant *Mtb* are exposed to *in vivo*, these differences are to be expected. Overall, the ability to compare proteomic regulation in *Mtb* across multiple dormancy models broadens our insight into the strengths of said strategies and challenges us to continue to develop better models for studying this deadly pathogen in the laboratory.

## Supporting information

Supplemental Tbl 1 and Fig S1

## Acknowledgements

Funding for this research was provided by the National Institute of Health (NIAID: R01 AI149737). Pacific Northwest National Laboratory is operated by Battelle for the Department of Energy (DOE) under contract DE-AC05-76RL01830. A portion of this research was supported by an Environmental Molecular Sciences Laboratory (EMSL) user project award (https://www.emsl.pnnl.gov/project/60231), for leveraging instrument capabilities operating at EMSL, a DOE Office of Science User Facility. We thank Priscila Lalli and Ron Moore for performing LC-MS analyses and Leo Gorham for help with sample preparation.

## Abbreviations

ACN: acetonitrile
AG: arabinogalactan
AGC: automatic gain control
Alr: alanine racemase
BCA: bicinchoninic acid
BPaL: bedaquiline, pertomanid, linezolid
CrfA: carbapenem resistance factor
CS: carbon starvation
Cyt *bd*: cytochrome *bd* oxidase
Ddl: D-alanine—D-alanine ligase
Ddn: deazaflavin-dependent nitroreductase
DOE: Department of Energy
FDR: false discovery rate
GarA: glycogen accumulation regulator A
GlcNAc: N-acetylglucosamine
HCD: higher-energy collisional dissociation
IMD: independence of missingness
K: lysine
LAM: lipoarabinomannan
LC-MS/MS: liquid chromatography tandem mass spectrometry
Ldt: L,D-transpeptidase
LM: lipomannan
LTBI: latent tuberculosis infection
MA: mycolic acids
MASIC: MS/MS automated selected ion chromatogram
*Mtb*: *Mycobacterium tuberculosis*
MurNAc: N-acetylmuramic acid
NDH-I: NADH dehydrogenase type I
NO: nitric oxide
OADC: Oleic Albumin Dextrose Catalase
P: pantothenate
PAV: peptide abundances vector
PG: peptidoglycan
PIM: phosphatidylinositol mannoside
Rep: replicating
rMd: robust Mahalanobis distance
SIC: selected ion chromatogram
SPE: solid phase extraction
STPK: serine/threonine protein kinase
TB: tuberculosis
TFA: trifluoroacetic acid
UPLC: ultra performance liquid chromatography

