## Supplemental Tbl 1 and Fig S1 for "Proteomic characterization of *Mycobacterium tuberculosis* subjected to carbon starvation"

**Table S1**. Culture growth under CS and Rep conditions for *M. tuberculosis* (mc^2^6020).

| **Culture** | **Starting OD_600_** | **Growth Time (d)** | **Harvest OD_600_** | **Lysate Conc (μg/mL)** |
| --- | --- | --- | --- | --- |
| CS- A | ~0.2 | 35 | 0.31 | 803 |
| CS- B | ~0.2 | 35 | 0.31 | 859 |
| CS- C | ~0.2 | 35 | 0.30 | 749 |
| CS- D | ~0.2 | 35 | 0.29 | 571 |
| CS- E | ~0.2 | 35 | 0.29 | 733 |
| CS- F | ~0.2 | 35 | 0.31 | 800 |
| Rep J | ~0.2 | 5 | 1.04 | 4178 |
| Rep K | ~0.2 | 5 | 0.93 | 3957 |
| Rep L | ~0.2 | 5 | 1.00 | 3812 |
| Rep M | ~0.2 | 5 | 0.92 | 2500 |
| Rep N | ~0.2 | 5 | 0.96 | 3423 |
| Rep O | ~0.2 | 5 | 1.10 | 3198 |

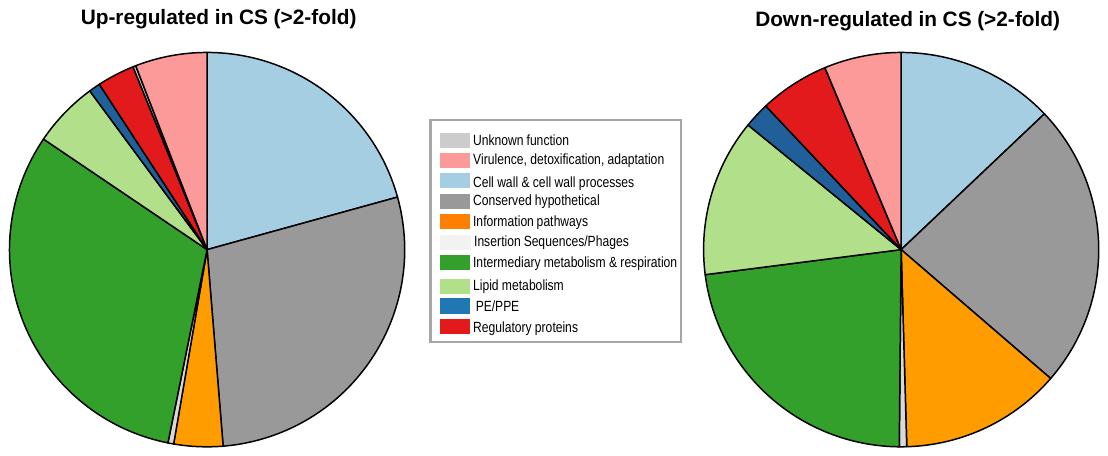

**Figure S1**. Functional classification of proteins differentially regulated under CS conditions.
